# Deciphering anomalous heterogeneous intracellular transport with neural networks

**DOI:** 10.1101/777615

**Authors:** Daniel S Han, Nickolay Korabel, Runze Chen, Mark Johnston, Viki J. Allan, Sergei Fedotov, Thomas A. Waigh

**Affiliations:** Department of Mathematics, University of Manchester, M13 9PL, UK; Department of Computer Science, University of Manchester, M13 9PL, UK; School of Biological Sciences, University of Manchester, M13 9PL, UK; Department of Physics and Astronomy, University of Manchester, M13 9PL, UK; The Photon Science Institute, University of Manchester M13 9PL, UK

## Abstract

Biological intracellular transport is predominantly heterogeneous in both time and space, exhibiting varying non-Brownian behaviour. Characterisation of this movement through averaging methods over an ensemble of trajectories or over the course of a single trajectory often fails to capture this heterogeneity adequately. Here, we have developed a deep learning feedforward neural network trained on fractional Brownian motion, which provides a novel, accurate and efficient characterization method for resolving heterogeneous behaviour of intracellular transport both in space and time. Importantly, the neural network requires significantly fewer data points compared to established methods, such as mean square displacements, rescaled range analysis and sequential range analysis. This enables robust estimation of Hurst exponents for very short time series data, making possible direct, dynamic segmentation and analysis of experimental tracks of rapidly moving cellular structures such as endosomes and lysosomes. By using this analysis, we were able to interpret anomalous intracellular dynamics as fractional Brownian motion with a stochastic Hurst exponent.

## Introduction

The majority of transport inside cells on the mesoscale (nm-100 μm) is now known to exhibit non-Brownian anomalous behaviour ***Metzler and Klafter (2004); Barkai et al. (2012); Waigh (2014)***. This has wide ranging implications for most of the biochemical reactions inside cells and thus cellular physiology. It is vitally important to be able to quantitatively characterise the dynamics of organelles and cellular responses to different biological conditions ***Van Bergeijk et al. (2015); Patwardhan et al. (2017); Moutaux et al. (2018)***. Classification of different non-Brownian dynamic behaviours at various time scales has been crucial to the analysis of intracellular dynamics ***Fedotov et al. (2018); Bressloff and Newby (2013)***, protein crowding in the cell ***Banks and Fradin (2005); Weiss et al. (2004a)***, microrheology ***Waigh (2005, 2016)***, entangled actin networks ***Amblard et al. (1996)***, and the movement of lysosomes ***Ba et al. (2018)*** and endosomes ***Flores-Rodriguez et al. (2011a)***. Anomalous transport is currently analysed by statistical averaging methods and this has been a barrier to understanding the nature of heterogeneous anomalous transport.

Spatiotemporal analysis of intracellular dynamics is often performed by acquiring and tracking microscopy movies of fluorescing membrane-bound organelles in a cell ***Rogers et al. (2007); Flores-Rodriguez et al. (2011a); Chenouard et al. (2014)***. These tracks are then commonly interpreted using statistical tools such as the mean square displacement (MSD) averaged over the ensemble of tracks, 〈Δ*r*^2^(*t*)〉. The MSD is a measure that is widely used in physics, chemistry and biology. In particular, MSDs serve to distinguish between anomalous and normal diffusion at different temporal scales by determining the anomalous exponent *α* through 〈Δ*r*^2^(*t*)〉 ~ *t^α^* ***Metzler and Klafter (2000)***. Diffusion is defined as *α* = 1, sub-diffusion 0 < *α* < 1 and super-diffusion 1 < *α* < 2 ***Klafter and Sokolov (2011)***. To improve the statistics of MSDs, they are often averaged over different temporal scales, forming the time-averaged MSD (TAMSD), 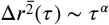, where *τ* is the lag time ***Sokolov (2012)***.

For stochastic processes with long-range time dependence such as fractional Brownian motion (fBm), other statistical averaging methods exist. For fBm, the MSD is 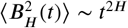 with the Hurst exponent, *H* varying between 0 and 1. One can use rescaled and sequential range analysis ***Samorodnitsky (2016); Peters (1994)*** to estimate *H*. The advantage of modelling intracellular transport with fBm is that both sub-diffusion (0 < *H* < 1/2) and super-diffusion (1/2 < *H* < 1) can be explained in a unified manner using only the Hurst exponent. The essence of fBm is that long-range correlations result in random trajectories that are anti-persistent (0 < *H* < 1/2) or persistent (1/2 < *H* < 1). The term persistence can be understood as the processive motor-protein transport of cargo in one direction, whether it be retrograde or anterograde. From a probabilistic viewpoint, persistence can be interpreted as the cargo being more likely to keep the same direction given it had been moving in this fashion before. On the other hand, how can we understand anti-persistence in intracellular transport? In the opposite sense, anti-persistence is interpreted as cargo being more likely to change its direction of movement given it had been moving in one before. Anti-persistence can arise if cargo is confined to a local volume in the cytoplasm simply due to crowding or tethering biochemical interactions ***Harrison et al. (2013)***, which in effect leads to sub-diffusion ***Weiss et al. (2004b); Ernst et al. (2012)***. By interpreting intracellular cargo transport as fBm, there are two main advantages: we can describe movement with the intuitive and biological concept of persistence and anti-persistence; and we can provide an immediate link to anomalous diffusion since *α* = 2*H* for constant *H*.

Cargo movement *in vivo* often exhibits random switching between persistent and anti-persistent movement, even in a single trajectory ***Chen et al. (2015)***. Therefore, we can model this by a stochastic local Hurst exponent, *H*(*t*), which jumps between persistent (1/2 < *H*(*t*) < 1) and anti-persistent (0 < *H*(*t*) < 1/2) states. Still, a major challenge exists: how can we estimate a local stochastic Hurst exponent from a trajectory?

Whilst exponent estimation using neural networks are emerging ***Bondarenko et al. (2016)***, segmentation of single trajectories into persistent and anti-persistent sections based on instantaneous dynamic behaviour has not been studied. Instead, hidden Markov models ***Monnier et al. (2015); Persson et al. (2013)*** and windowed analyses ***Getz and Saltz (2008)*** are commonly used to segment local behaviour along single trajectories. Even so, most methods neglect the microscopic processes which are often a feature of intracellular transport (e.g. alternation between “runs” and “rests”) ***Weiss et al. (2004a); Chen et al. (2015); Fedotov et al. (2018)*** and the non-Markovian nature of their motion ***Fuliński (2017)***.

Here, we present a new method for characterising anomalous transport inside cells based on a Deep Learning Feedforward Neural Network (DLFNN) that is trained on fBm. Neural networks are becoming a general tool in a wide range of fields, such as single-cell transcriptomics 32 and protein folding 33. We find the neural network is a much more sensitive method to characterise fBm than previous statistical tools, since it is an intrinsically non-linear regression method that accounts for correlated time series. In addition, it can estimate the Hurst exponent using as few as 7 consecutive time points with good accuracy. The new method enables the interpretation of experimental trajectories of lysosomes and endosomes as fBm with stochastic local Hurst exponent, *H*(*t*). This in turn allows us to unambiguously and directly classify endosomes and lysosomes to be in anti-persistent or persistent states of motion at different times. From experiments, we observe that the time spent within these two states both exhibit truncated heavy-tailed distributions.

To our knowledge, this is the first method which is capable of resolving heterogeneous behaviour of anomalous transport in both time and space. We anticipate that this method will be useful in characterising a wide range of systems that exhibit anomalous heterogeneous transport. We have therefore created a GUI computer application called DeepExponent in which the DLFNN is implemented so that the wider community can conveniently access this analysis method

## Results and Discussion

### The DLFNN is more accurate than established methods

We tested a DLFNN trained on fBm with 3 hidden layers of densely connected nodes on *N* = 10^4^ computer-generated fBm trajectories each with *n* = 10^2^ evenly spaced time points and constant Hurst exponent *H_sim_*, randomly chosen between 0 and 1. The DLFNN estimated the Hurst exponents *H_est_* based on the trajectories, and these were compared with those estimated from TAMSD, rescaled range, and sequential range methods (Figure 1a). The difference between the simulated and estimated values Δ*H* = *H_sim_* – *H_est_* was much smaller for the DLFNN than for the other methods (Figure 1a), and the DLFNN was ~ 3 times more accurate at estimating Hurst exponents with a mean absolute error (*σ_H_*) ~ 0.05. Also, the errors in estimation of the DLFNN are more stable across values of *H_sim_*.

**Figure 1.**
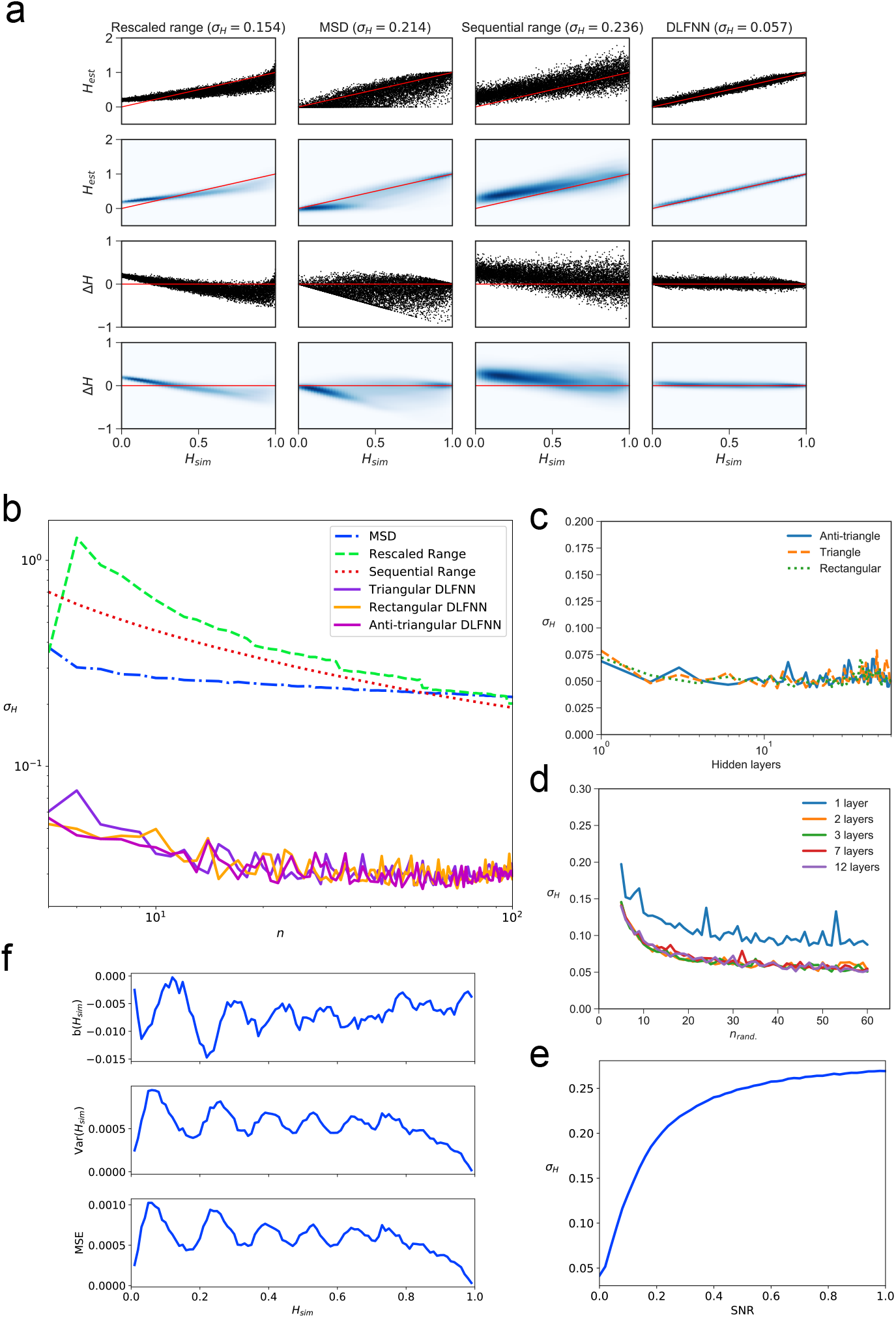
Plots showing tests of exponent estimation for the DLFNN using *N* = 10^4^ simulated fBm trajectories. **a**) Plots showing the Hurst exponent estimates of fBm trajectories with *n* = 10^2^ data points by triangular DLFNN with three hidden layers compared with conventional methods. Plots are vertically grouped by Hurst exponent estimation method: (left to right) rescaled range, MSD, sequential range and DLFNN. *σ_H_* values are shown in the title. *Top row*: Scatter plots of estimated Hurst exponents *H_est_* and the true value of Hurst exponents from simulation *H_sim_*. The red line shows perfect estimation. *Second row*: Due to the density of points, a gaussian kernel density estimation was made of the plots in the top row (see methods). *Third row*: Scatter plots of the difference between the true value of Hurst exponents from simulation and estimated Hurst exponent Δ*H* = *H_sim_* – *H_est_*. *Last row*: Gaussian kernel density estimation of the plots in the third row. **b**) *σ_H_* as a function of the number of consecutive fBm trajectory data points *n* for different methods of exponent estimation. **c**) *σ_H_* as a function of the number of hidden layers in the DLFNN for rectangular, triangular and anti-triangular structures (see methods). **d**) *σ_H_* as a function of the number of randomly sampled fBm trajectory data points *n_rand_* with different number of hidden layers in the DLFNN shown in the legend. **e**) *σ_H_* as a function of the signal-to-noise ratio (SNR) from Gaussian random numbers added to all *n* = 10^2^ data points in simulated fBm trajectories. **f**) Plots of bias *b*(*H_sim_*), variance Var(*H_sim_*) and mean square error (MSE) as functions of *H_sim_*. For each value of *H_sim_*, fBm trajectories with *n* = 100 points were simulated and estimated by a triangular DLFNN.

Tracking of intracellular motion usually generates trajectories with a variable number of data points. We therefore compared the performance of the different exponent estimation methods when the number of evenly spaced, consecutive fBm time points in a trajectory varied between *n* = 5,6,…, 10^2^ points. The DLFNN maintained an accuracy of *σ_H_* ~ 0.05 across *n*, whereas *σ_H_* of other methods increase as *n* decreases (Figure 1b), and was always substantially worse than that of the DLFNN estimation. Different topologies of DLFNN structure (see methods Figures 7, 8 and 9) performed similarly (Figure 1b and c), and introducing more hidden layers did not affect the accuracy of estimation (Figure 1c). Given that the structure of DLFNN does not significantly affect the accuracy of exponent estimation, a triangular densely connected DLFNN was used for all subsequent analyses.

The structure of a triangular DLFNN means that the input layer consists of *n* nodes, which are densely connected to *n* – *l* nodes in the first hidden layer, such that at the *l*^th^ hidden layer, there would be *n* – *l* densely connected nodes. Then to estimate the Hurst exponent these nodes are connected to a single node using a Rectified Linear Unit(ReLU) activation function, which returns the exponent estimate. A triangular DLFNN therefore uses only 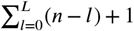 nodes for *L* hidden layers and *n* input points, whereas the rectangular structure uses *nL* + 1 nodes and the anti-triangular uses 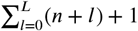. The triangular structure results in a significant decrease in training parameters, and hence computational requirements, while maintaining good levels of accuracy. This demonstrates that a computationally inexpensive neural network can accurately estimate exponents.

The DLFNN’s estimation capabilities were tested further by inputting *n_rand_* randomly sampled time points from the original fBm trajectories. Surprisingly, *σ_H_* ~ 0.05 is regained even with just 40 out of 100 data points randomly sampled from the time series for any triangular DLFNN with more than 1 hidden layer (Figure 1d). For this method to work with experimental systems, it must estimate Hurst exponents even when the trajectories are noisy. Figure 1e shows how the exponent estimation error increases when Gaussian noise with increasing signal-to-noise ratio (SNR) is added to the fBm trajectories. Importantly, the DLFNN accuracy *σ_H_* at 20% SNR is as good as the accuracy of other methods with no noise (compare Figure 1a and e).

To characterise the accuracy of *H_sim_* estimation by the DLFNN, we calculated the bias, 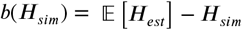; variance, 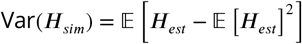; and mean square error, MSE = Var(*H_sim_*) + *b*(*H_sim_*)^2^ (Figure 1f). To quantify the efficiency of the estimator the Fisher information of the neural network’s estimation needs to be found and the Cramer-Rao lower bound calculated. The values of bias, variance and MSE were very low (Figure 1f), which taken together with the simplicity of calculation and the accuracy of estimation even with small number of data points, demonstrates the strength of the DLFNN method. Furthermore, once trained, the model can be saved and reloaded at any time. Saved models and software available for download in the Supplementary Materials.

### DLFNN allows analysis of trajectories with local stochastic Hurst exponents

Estimating local Hurst exponents is fundamentally important because much research has focused on inferring active and passive states of transport within living cells using position-derived quantities such as windowed MSDs, directionality and velocity ***Arcizet et al. (2008); Monnier et al. (2015)***. The trajectories are then segmented and Hurst exponents measured in an effort to characterise the behaviour of different cargo when they are actively transported by motor proteins ***Chen etal. (2015); Fedotov et al. (2018)*** or sub-diffusing in the cytoplasm ***Jeon et al. (2011)***. However, conventional methods such as the MSD and TAMSD need trajectories with many time points (*n* ~ 10^2^ – 10^3^) to calculate a single Hurst exponent value with high fidelity. In contrast, the DLFNN enables the Hurst exponent to be estimated, directly from positional data, for a small number of points. Furthermore, the DLFNN measures local Hurst exponents without averaging over time points and is able to characterize particle trajectories that may exhibit multi-fractional, heterogeneous dynamics.

To provide a synthetic data set that mimics particle motion in cells, we simulated fBm trajectories with Hurst exponents that varied in time, and applied a symmetric moving window to estimate the Hurst exponent using a small number of data points before and after each time point (Figure 2). The DLFNN was able to identify segments with different exponents, and provided a good running estimation of the Hurst exponent values. The DLFNN could also handle trajectories with different diffusion coefficients, and generally performed better than MSD analysis when a sliding window was used (see Appendix A).

**Figure 2.**
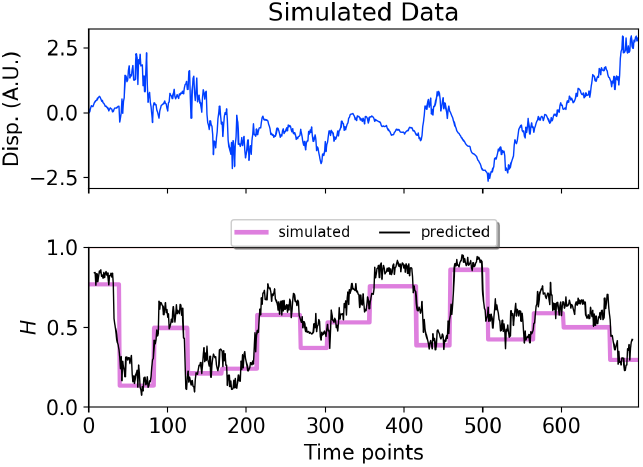
*Top*: Plot of displacement as a function of time from a simulated fBm trajectory (blue) with multiple exponent values. *Bottom*: Hurst exponent values used for simulation (magenta), and the DLFNN exponent predictions of the neural network using a 14 point moving window (black). The red line marks diffusion, *H* = 0.5.

We then applied this method to an experimental trajectory obtained from automated tracking ***Newby et al. (2018)*** data of GFP-Rab5 labelled endosomes in a living MRC-5 cell (Figure 3). A moving window of 14 points identified persistent (green) and anti-persistent (magenta) segments, which corresponded well to the moving window velocity plots (Figure 3, lower panel), confirming that the neural network is indeed distinguishing passive states from active transport states with non-zero average velocity.

**Figure 3.**
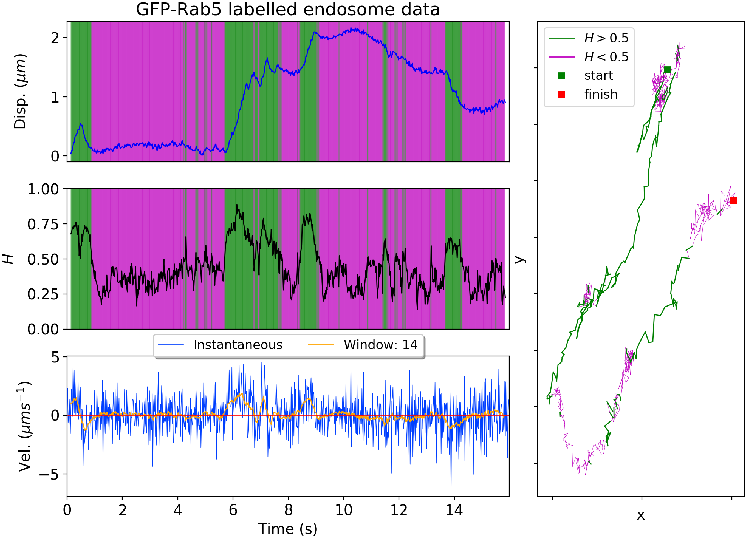
*Top*: Plot of displacement from a single GFP-Rab5 expressing endosome trajectory in an MRC-5 cell (blue). Shaded areas show persistent (0.5 < *H* < 1 in green) and anti-persistent (0 < *H* < 0.5 in magenta) behaviour. *Middle*: A 14 point moving window DLFNN exponent estimate for the trajectory (black) with a line (dashed) marking diffusion *H* = 0.5. *Bottom*: Plot of instantaneous and moving (14 point) window velocity. *Right*: Plot of the trajectory of a GFP-Rab5 expressing endosome in an MRC-5 cell with start and finish positions, and, persistent (green) and anti-persistent (magenta) segments.

### Trajectory analysis using DLFNN shows regime switching

To understand organelle motility in the context of cell behaviour, an additional layer of complexity needs to be considered – the location of the moving structure within the cell itself. Such information would reveal zones that favour anti-persistent or persistent movement ***Bálint et al. (2013)***. Using the neural network, trajectories of GFP-Rab5 endosomes from a single MRC-5 cell were plotted with colours depicting the changing Hurst exponent at different points in each trajectory (Figure 4), which ranged from 0 to 1. This revealed an enrichment of anti-persistent organelles in the cell periphery, with long-range persistent movement towards the nucleus occuring within more central regions, as expected ***Flores-Rodriguez et al. (2011b)***.

**Figure 4.**
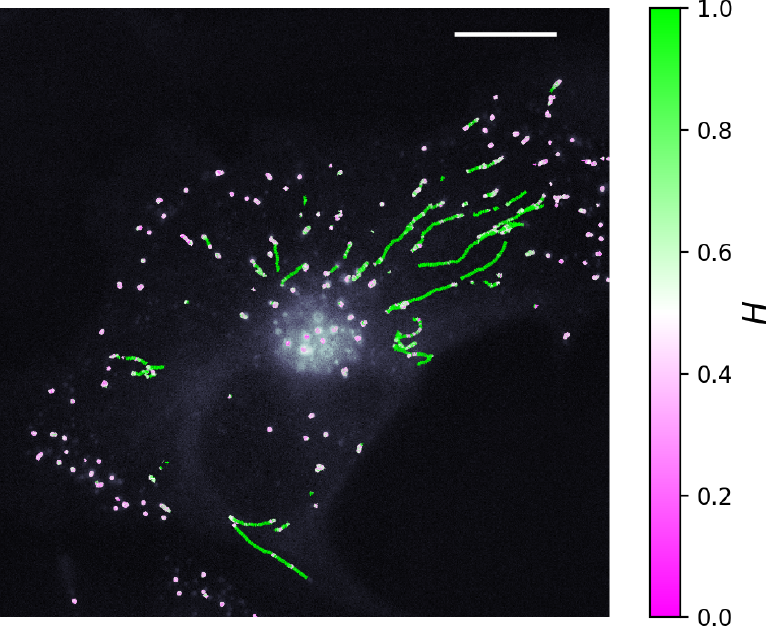
An MRC-5 cell stably expressing GFP-Rab5, with tracking data overlaid. The colours show the value of *H* estimated by the neural network using a 14 point window. The scalebar is 10 μm

The accurate estimation of Hurst exponents from very few data points also allowed us to analyse intracellular dynamics by segmenting individual endosomal and lysosomal trajectories into persistent and anti-persistent sections. A total of 65 MRC5 cells for endosomes and 71 MRC5 cells for lysosomes were analysed, giving 40,800 (endosome) and 38,039 (lysosome) tracks that were segmented into 277,926 (endosome) and 474,473 (lysosome) sections, each yielding a displacement, duration and average *H*.

A histogram of Hurst exponent was plotted, and by fitting a six component Gaussian mixture model to this distribution, individual segments were automatically binned into the appropriate Gaussian distribution according to their Hurst exponent (Figure 5a). Then, plots of the displacements, average velocities and duration of these segments were made to check that the overall average Hurst exponents of each Gaussian distribution agreed with the general properties of the segments (Figures 5b and c). The segments classed with *H* > 0.5 had linear correlations between displacements and time, which was not seen for segments with *H* < 0.5. The anti-persistent segments generally had a longer duration than the persistent segments, but had much smaller displacements (note the different axes scales between upper and lower rows in Figures 5b and c). This is expected because lysosomes and endosomes are known to spend the majority of time in a subdiffusive state interrupted by brief periods of active motor transport ***Flores-Rodriguez et al. (2011a)***.

**Figure 5.**
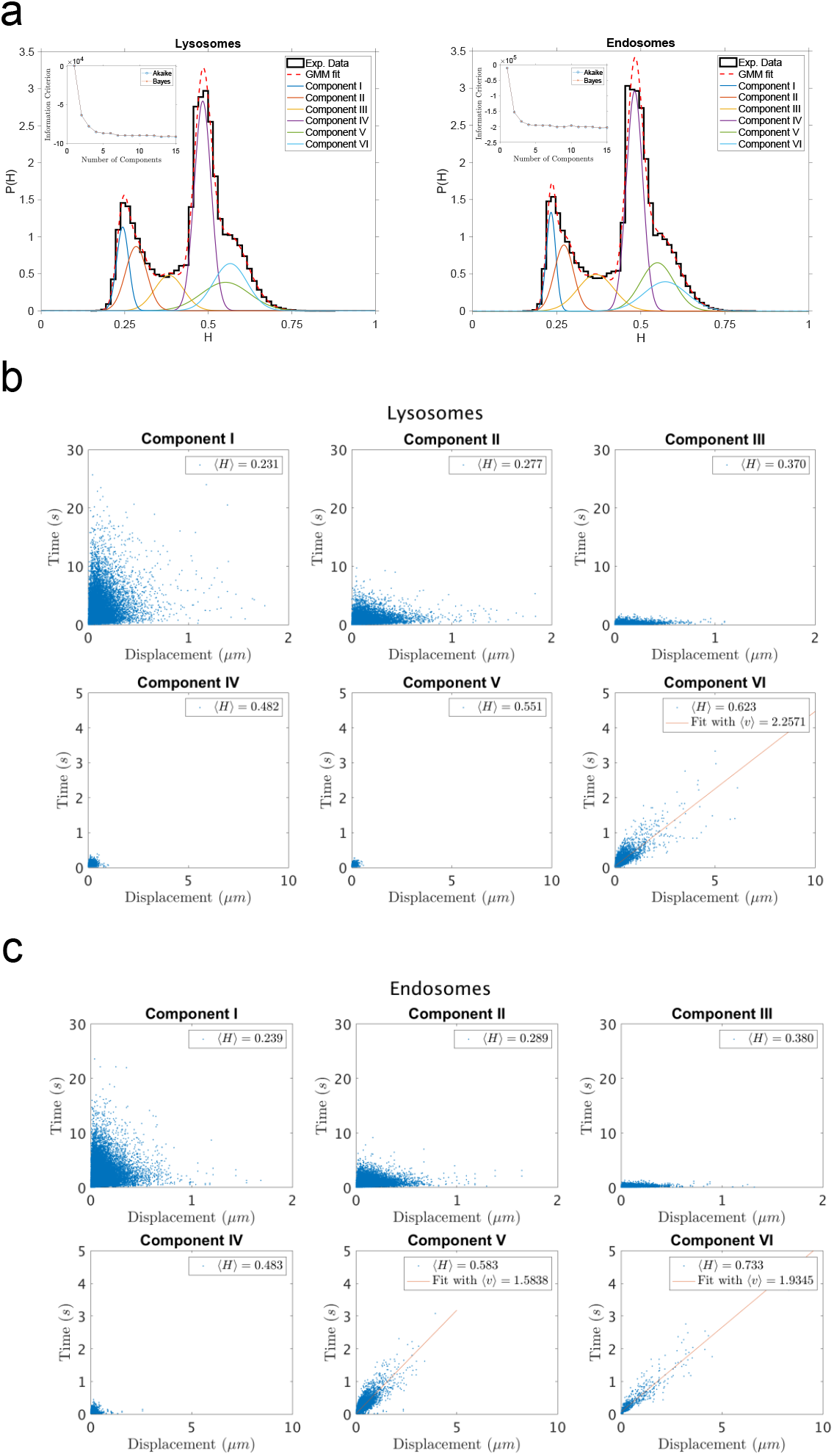
**a**) Histograms of Hurst exponents (black) and the Gaussian mixture model fit for 6 components (red) for lysosomes and endosomes. The individual Gaussian distributions are shown on the same plot. The number of components were chosen through the Bayes information criterion. *Inset*: The Akaike and Bayes information criterion against number of components in the Gaussian mixture model. **b**) Scatter plot of segment displacement and segment duration time for lysosome trajectories grouped in different clusters by the Gaussian mixture model. The legend shows the mean Hurst exponent, *H*, for each cluster. **c**) The same analysis as shown in **b** for endosome trajectories.

Here, we have not used any additional assumptions to extract these states from the data. The different anti-persistent and persistent states arise naturally out of the trajectories due to the local Hurst exponent estimates. An intriguing possibility is that the mixture of Gaussian distributions in Figure 4 reflects separate transient biochemical states of lysosomes and endosomes that correlate with their motility. The DLFNN is therefore a powerful tool, since the change in the distribution of Hurst exponent for different biological conditions can be quantified directly from the motility of intracellular organelles or any object that can be tracked.

### fBm with a stochastic Hurst exponent is a new intracellular transport model

fBm is a Gaussian process *B_H_*(*t*) with zero mean and covariance 〈*B_H_*(*t*)*B_H_*(*s*)〉 ~ *t*^2*H*^ + *s*^2*H*^ – (*t* – *s*)^2*H*^, where the Hurst exponent, *H* is a constant between 0 and 1. With the DLFNN providing local estimates of the Hurst exponent, we discovered that the motion of endosomes and lysosomes can be described as fBm with a stochastic Hurst exponent, *H*(*t*). This is different to multifractional Brownian motion ***Peltier and Véhel (1995)*** where *H*(*t*) is a function of time. In our case, *H*(*t*) is itself a stochastic process. Furthermore, Figure 3 shows the Hurst exponent varying between persistent and anti-persistent states indicating that *B_H_*(*t*) exhibits regime switching behaviour.

We found that the times that lysosomes and endosomes spend in a persistent and anti-persistent state are heavy-tailed (Figure 6). These times are characterised by the probability densities *ψ*(*t*) ~ *t*^−*μ*−1^, where anti-persistent states have 0 < *μ* < 1 and persistent states have 1 < *μ* < 2. Extensive plots and fittings are shown in Figure 6 and Supplementary Material B. In fact, the residence time probability density has an infinite mean to remain in an anti-persistent state (0 < *H*(*t*) < 1/2) but in persistent states (1/2 < *H*(*t*) < 1) the mean of the residence time probability density is finite and the second moment is infinite. This implies that the vesicles may have a biological mechanism to prioritise certain interactions within the complex cytoplasm, similar to how human dynamics are often heavy tailed and bursty ***Barabasi (2005)***.

**Figure 6.**
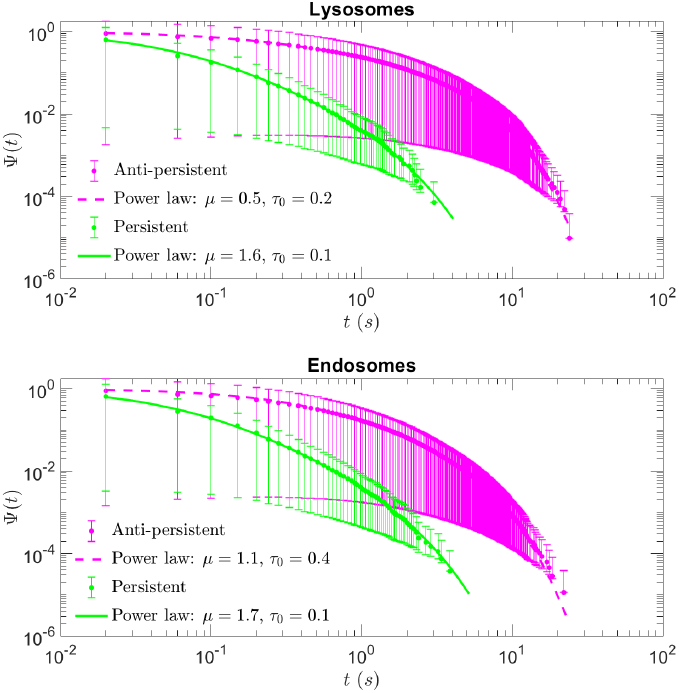
A plot of survival functions with error bars for persistent and anti-persistent segments for lysosomes and endosomes with the power-law fits. Fit parameters can be found in Table 1

## Conclusions

We developed a Deep Learning Feedforward Neural Network trained on fBm that estimates accurately the Hurst exponent for heterogeneous trajectories. Estimating the Hurst exponent using a DLFNN is not only more accurate than conventional methods but also enables direct trajectory segmentation without a drastic increase in computational cost. We package this DLFNN analysis code into a user-friendly application called DeepExponent, which can predict the Hurst exponent with consistent accuracy for as few as 7 consecutive data points. This is useful to biologists since major limitations to trajectory analysis are: the brevity of tracks due to the fact that particles may rapidly switch between motile states or move out of the plane of focus;the rapid nature of some biochemical reactions;and the bleaching of fluorescent probes (with non-bleaching probes often being bulky or cytotoxic). This method can be used to detect persistent and anti-persistent states of motion purely from the positional data of trajectories and removes the prerequisite of time or ensemble averaging for effective heterogeneous transport characterisation. The DLFNN enabled us to discover regime switching in lysosome and endosome movement that can be modelled by fBm with a stochastic Hurst exponent. This interpretation is a unified approach to describing motion with anti-persistence and persistence varying over time. Furthermore, the residence time of vesicles in a persistent or anti-persistent state is found to be heavy tailed, which implies that endosomes and lysosomes possess biological mechanisms to prioritise varying biological processes similar to human dynamics ***Barabasi (2005)***. We hope that this type of analysis will allow discoveries in particle motility of a more refined nature and make applying anomalous transport theory more accessible to researchers in a wide variety of disciplines.

## Methods and Materials

### Hurst exponent estimation methods

**Time averaged MSDs** were calculated using

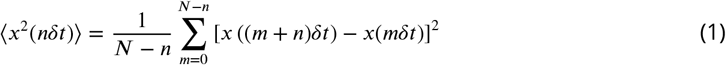

where *x*(*nδt*) is the track displacement at time *nδt* and a track contains *N* coordinates spaced at regular time intervals of *δt*. From now on, 〈*x*〉 will denote the time average of *x* unless explicitly specified otherwise. The total time is *T* = (*N* – 1)*δt* and *n* = 1,2, …, *N* – 1. Lag-times are the set of possible *nδt* within the data set and 〈*x*^2^(*nδt*)〉 was then fit to a power-law ~ *t*^2*H*^ using the ‘scipy.optimize’ package in Python3 to estimate the exponent *H*.

**Rescaled ranges** were calculated by creating a mean adjusted cumulative deviate series 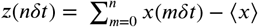 from original displacements *x*(*nδt*) and mean displacement 〈*x*〉. Then the rescaled range is calculated by

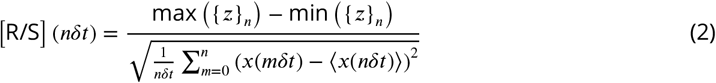

where {*z*}_*n*_ = *z*(0),*z*(*δt*),*z*(2*δt*),…, *z*(*nδt*). The rescaled range is then fitted to a power law [R/S] (*nδt*) ~ (*nδt*)^*H*^ where *H* is the Hurst exponent ***Hurst (1951)***. The ‘compute_Hc’ function in the ‘hurst’ package in Python3 estimates the Hurst exponent in this way.

**Sequential ranges** are defined as

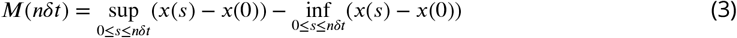

where sup(*x*) is the supremum and inf(*x*) is the infimum for the set *x* of real numbers. Then *M*(*nδt*) = (*nδt*)^*H*^ *M*(*δt*) ***Feller (1951)***.

### DLFNN structure and training

The fractional Brownian trajectories were generated using the Hosking method within the “FBM” function available from the ‘fbm’ package in Python3. The DLFNN was built using Tensor-flow ***Abadi et al. (2016)*** and Keras ***Chollet (2015)*** in Python3 and trained by using the simulated fractional Brownian trajectories. The training and testing of the neural network were performed on a workstation PC equipped with 2 CPUs with 32 cores (Intel(R) Xeon CPU E5-2640 v3) and 1 GPU (NVIDIA Tesla V100 with 16 GB memory). The structure of the neural network was a multilayer, feedforward neural network where all nodes of the previous layer were densely connected to nodes of the next layer. Each node had a ReLU activation function and the parameters were optimized using the RMSprop optimizer (see Keras documentation ***Chollet (2015)***). Three separate structures were explored and examples of these structures for 2 hidden layers and 5 time point inputs are shown in Figure 7, 8 and 9. The triangular structure was predominantly used since this was the least computationally expensive and accuracy between different structures were similar. To compare the accuracy of different methods, the mean absolute error (*σ_H_*) of *N* trajectories, 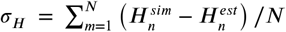, was used. Before inputting values into the neural network, the time series was differenced to make it stationary. The input values of a fBm trajectory {*x*} = *x*_0_,*x*_1_,…,*x_n_* were differenced and normalized so that {*x_input_*} = (*x*_1_ – *x*_0_)/range(*x*), (*x*_2_ – *x*_1_)/range(*x*),…, (*x_n_* – *x*_*n*−1_)/range(*x*).

**Figure 7.**
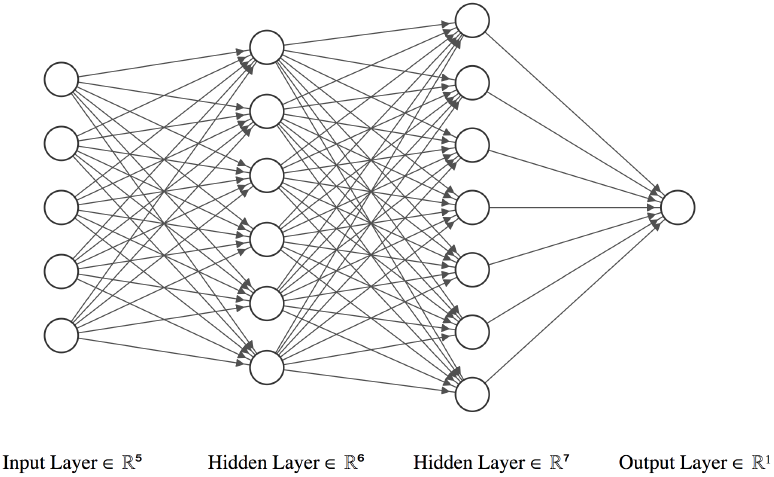
An ‘anti-triangular’ structure example for 2 hidden layers and *n* = 5 time series input points.

**Figure 8.**
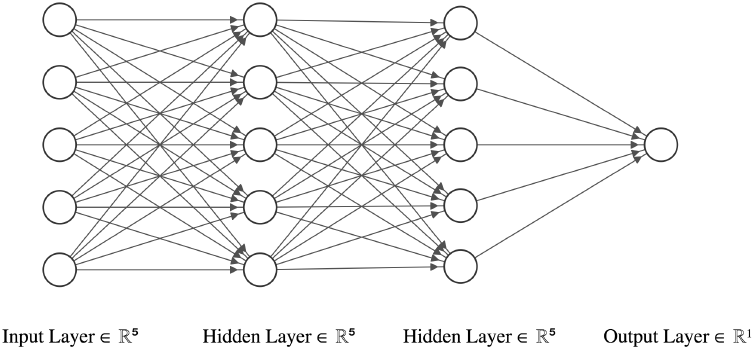
An ‘rectangular’ structure example for 2 hidden layers and *n* = 5 time series input points.

**Figure 9.**
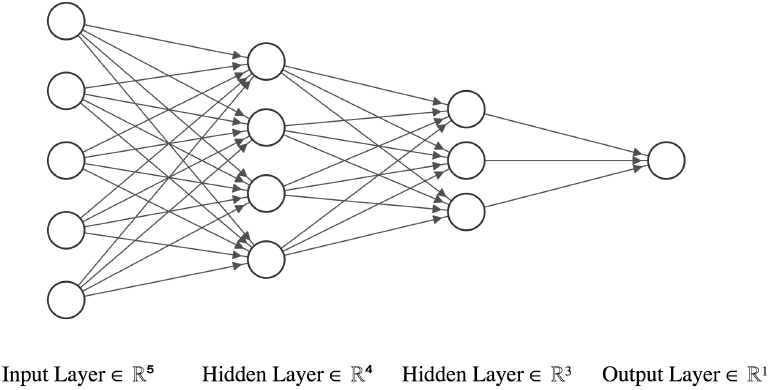
An ‘triangular’ structure example for 2 hidden layers and *n* = 5 time series input points.

### Gaussian kernel density estimation

Kernel density estimation (KDE) is a non-parametric method to estimate the probability density function (PDF) of random variables. If *N* random variables *x_n_* are distributed by an unknown density function *P*(*x*), then the kernel density estimate *P*(*x*) is

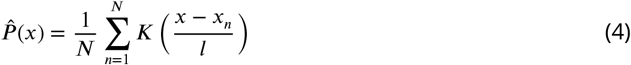

where *K*(·) is the kernel function and *l* is the bandwidth. In this paper, we have used a Gaussian KDE, 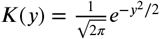, to estimate the two dimensional PDFs of the second and bottom row in Figure 1a. This was performed in Python3 using ‘scipy.stats.gaussian_kde’ and Scott’s rule of thumb for bandwidth selection.

### Biological experiments and data acquisition

An MRC5 cell line stably expressing GFP-Rab5C was kindly provided by Drs. Guy Pearson and Evan Reid (Cambridge Institute for Medical Research, University of Cambridge). To generate this cell line, GFP-Rab5C was PCRed from pIRES GFP-Rab5C (Seaman, M.; 2004) using Hpa1 GFP Forward’ (TAGGGAGTTAACATGGTGAGCAAGGGCGAGGA) and ‘Not1 Rab5C Reverse’ (ATCCCTGCG-GCCGCTCAGTTGCTGCAGCACTGGC) oligonucleotide primers. The GFP-Rab5C PCR product and a pLXIN-I-NeoR plasmid were digested using Hpa1 (New England Biolabs – R0105) and Not1 (New England Biolabs – R3189) restriction enzymes. The GFP-Rab5C PCR product was then ligated into the digested pLXIN-I-NeoR using T4 DNA Ligase (New England Biolabs – M0202). The ligated plasmid was amplified in bacteria selected with ampicillin and verified using Sanger Sequencing. To generate the GFP-Rab5C MRC5 cell line, Phoenix retrovirus producer HEK293T cells were transfected with the pLXIN-GFP-Rab5C-I-NeoR plasmid to generate retrovirus containing GFP-Rab5C. MRC5 cells were inoculated with the virus, and successfully transduced cells were selected using 200 μm/mL Geneticin (G418 – Sigma-Aldrich G1397). Cells used were not clonally selected.

Stably expressing MRC5 cells were co-stained with LysoBrite (AAT Bioquest) were imaged using fluorescence microscopy and tracked with NNT (aitracker.net ***Newby et al. (2018)***). The cells were grown in MEM (Sigma Life Science) and 10% FBS (HyClone) and incubated for 48 hrs at 37°C in 5% CO_2_ on 35 mm glass-bottomed dishes (*μ*-Dish, Ibidi, Cat. No. 81150). For LysoBrite staining, LysoBrite was diluted 1 in 500 with Hanks Balanced Salt solution (Sigma Life Science). Then 0.5 mL of this solution was added to a 35 mm dish containing 2 mL of growing media with cells. Once at least 1 hr has passed, the cells were washed with PBS (Sigma Life Science) and the media is replaced with growing media.

After at least 6 hrs incubation, the growing media is replaced with live-imaging media composed of Hanks Balanced Salt solution (Sigma Life Science, Cat. No. H8264) with added essential and non-essential amino acids, glutamine, penicillin/streptomycin, 25mM HEPES (pH7.0) and 10% FBS (HyClone). The live-cell imaging was performed on an inverted Olympus IX71 microscope with an Olympus 100×1.35 oil PH3 objective. A Prime 95B sCMOS Camera (Photometrics) and OptoLED (Cairn Research) light source was used for the continuous imaging of MRC-5 cells. Filter sets ET470/40x and ET573/35x (Chroma Technology Corp.) were used for imaging GFP-Rab5 endosomes and Lysobrite stained lysosomes respectively. Endosomes and lysosomes were imaged in separate experiments. A stream of 20ms exposure was collected for 17 seconds using Metamorph software while the cells were kept at 37 °C (in atmospheric CO_2_ levels).

## Acknowledgments

The authors would like to thank: Dr. Jay Newby and Zach Richardson (UNC Chapel Hill) for assistance in using the automated tracking system (AITracker); Prof. Matthias Weiss (Universität Bayreuth), Dr. Henry Cox, Jack Hart and Hannah Perkins for discussions; and, Dr. Guy Pearson and Dr. Evan Reid (University of Cambridge) for providing the GFP-Rab5 MRC5 cell line. DH acknowledges financial support from the Wellcome Trust Grant No. 215189/Z/19/Z. SF, NK, TAW, and VJA acknowledge financial support from EPSRC Grant No. EP/J019526/1.

## Appendix 1

### Testing DLFNN accuracy for different diffusion coefficients

The DLFNN was compared to the MSD estimation method for trajectories with different diffusion coefficients to ensure that the DLFNN estimation was not scale dependent.

**Appendix 1 Figure 1.**
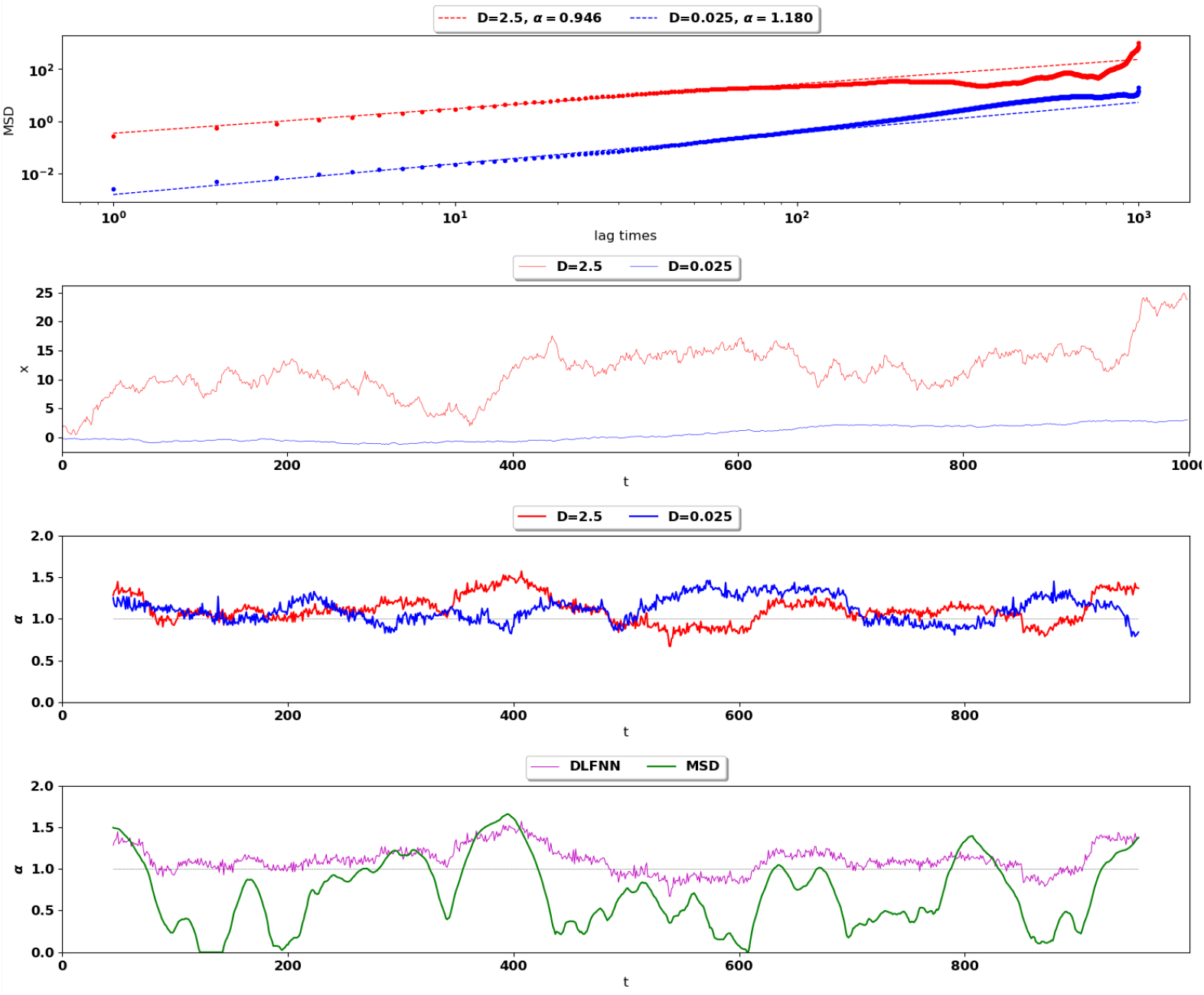
Top: MSD(points)and power-lawfits(dashed)fortwo different Brownian trajectories containing 1000 data points with diffusion coefficient 2.5 (red) and 0.025 (blue). The Hurst exponent should be *α* = 2*H* = 1 for both trajectories. Second Row: Simulation ofthe two Brownian trajectorieswith diffusion co-efficient 2.5 (red) and 0.025 (blue) Third Row: Local Hurst exponent estimates given by the DLFNN for the two different trajectories using a 90 point window. The averages of DLFNN Hurst exponent estimates are *α* = 2*H* = 1.110 (red) and *α* = 2*H* = 1.114 (blue). Bottom: Local Hurst exponent estimates given by DLFNN and MSDs using a 90 point window. The average of DLFNN Hurst exponent estimates is *α* = 2*H* = 1.110 and the average of MSD estimates is *α* = 2*H* = 0.937.

### Measuring the residence time and flight length probability density functions of persistent and anti-persistent states

Classifying persistent and anti-persistent states by Hurst exponent values 1/2 < *H* < 1 and 0 < *H* < 0.5 respectively, individual lysosome and endosome trajectories were segmented with a moving window of 9 points. Data was extracted from microscopy movies of 120 MRC5 cells acquired from three independent experiments. Endosomes from 60 cells and lysosomes from 60 different cells were tracked using AITracker ***Newby et al. (2018)***. Then the trajectories were segmented into anti-persistent (0 < *H* < 0.5) and persistent (0.5 < *H* < 1) using the Hurst exponent estimates by DLFNN. The time duration and particle displacement of these segments were measured and then fitted to distributions. In this way, we could measure the stochastic switching between active and passive transport and the statistics of vesicle movement within these states.

Figure 2 shows the survival time probabilities Ψ(*t*) of different states of motion (persistent and anti-persistent) in vesicle trajectories. Ψ(*t*) is the probability that the vesicle will still be in the same state of motion after time *t* has elapsed. Figure 2 shows that the persistent and anti-persistent states follow 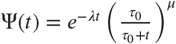.

The survival time probabilities show that lysosomes and endosomes are far more likely to remain trapped in a anti-persistent state than be persistently transported by motor proteins. While this is intuitively obvious in the context of cell biology, this analysis provides quantitative characterization of endosomal and lysosomal motility. Table 1 shows the full details of the fit results to the survival probabilities shown in Fig. 2. Figure 3 shows the empirical probability density functions (PDF) of the particle displacements for different states of motion. Displacements of segments are fitted to Burr Type XII distributions. Table 2 shows the full details of parameter estimates of Figure 3.

As expected, this analysis demonstrates that both endosomes and lysosomes are far more likely to move large distances when they are in the persistent state. This reconciles how vesicles are able to move large distances even though they are more likely to stay in a anti-persistent state for long periods of time. The large displacements in the persistent state compete with long durations spent in the anti-persistent state.

**Appendix 1 Figure 2.**
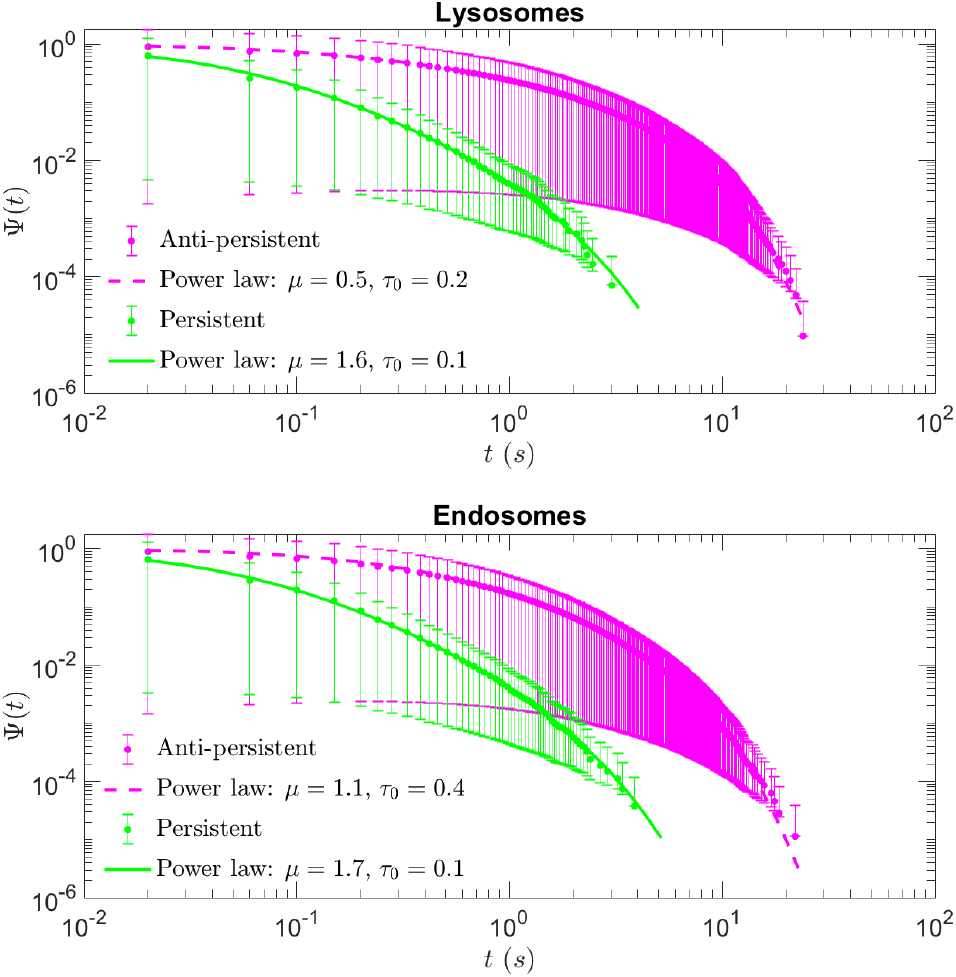
Plots showing the Kaplan-Meier estimates of survival fucntions Ψ(*t*) from lysosome and endosome experimental trajectories segmented using DLFNN (markers) and the corresponding fits (lines) for separate behaviours in vesicle trajectories. Error bars show 95% upper and lower confidence bounds calculated using Greenwood’s formula. Further details on the functional form of Ψ(*t*) and parameters are shown in Table 1.

**Appendix 1 Table 1.** Results for the fits of survival time probabilities shown in Fig 1. The parameters and the analytical survival functions used to fit the Kaplan-Meier estimator survival curves.

| Lysosome data | Survival function $\Psi(t)$ | Fit parameters for Fig. 2 |
| --- | --- | --- |
| Anti-persistent | $e^{-\lambda t} \left( \frac{\tau_0}{\tau_0 + t} \right)^\mu$ | $\mu = 0.54 \pm 0.005, \tau_0 = 0.151 \pm 0.002s, \lambda = 0.341 \pm 0.002s^{-1}$ |
| Persistent | $e^{-\lambda t} \left( \frac{\tau_0}{\tau_0 + t} \right)^\mu$ | $\mu = 1.63 \pm 0.10, \tau_0 = 0.062 \pm 0.006s, \lambda = 0.900 \pm 0.128s^{-1}$ |
| Endosome data | Survival function $\Psi(t)$ | Fit parameters for Fig. 2 |
| Anti-persistent | $e^{-\lambda t} \left( \frac{\tau_0}{\tau_0 + t} \right)^\mu$ | $\mu = 1.086 \pm 0.014, \tau_0 = 0.370 \pm 0.007s, \lambda = 0.360 \pm 0.005s^{-1}$ |
| Persistent | $e^{-\lambda t} \left( \frac{\tau_0}{\tau_0 + t} \right)^\mu$ | $\mu = 1.718 \pm 0.065, \tau_0 = 0.067 \pm 0.004s, \lambda = 0.768 \pm 0.083s^{-1}$ |

**Appendix 1 Table 2.** Results for the fits of displacement probability density functions shown in Fig 3. Displacements of segments in Fig 3 are fitted to Burr Type XII distributions,

| Lysosome data | PDF $P(x)$ | Fit parameters for Fig. 3 |
| --- | --- | --- |
| Anti-persistent | Burr Type XII | $c = 1.44 \pm 0.01, k = 1.93 \pm 0.05, x_0 = 0.063 \pm 0.002\mu m$ |
| Persistent | Burr Type XII | $c = 1.52 \pm 0.02, k = 0.79 \pm 0.02, x_0 = 0.023 \pm 0.001\mu m$ |
| Endosome data | PDF $P(x)$ | Fit parameters for Fig. 3 |
| Anti-persistent | Burr Type XII | $c = 1.49 \pm 0.01, k = 1.94 \pm 0.06, x_0 = 0.104 \pm 0.004\mu m$ |
| Persistent | Burr Type XII | $c = 1.62 \pm 0.03, k = 0.79 \pm 0.03, x_0 = 0.039 \pm 0.001\mu m$ |

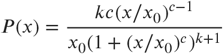

where *x, x*_0_, *c* and *k* > 0.

**Appendix 1 Figure 3.**
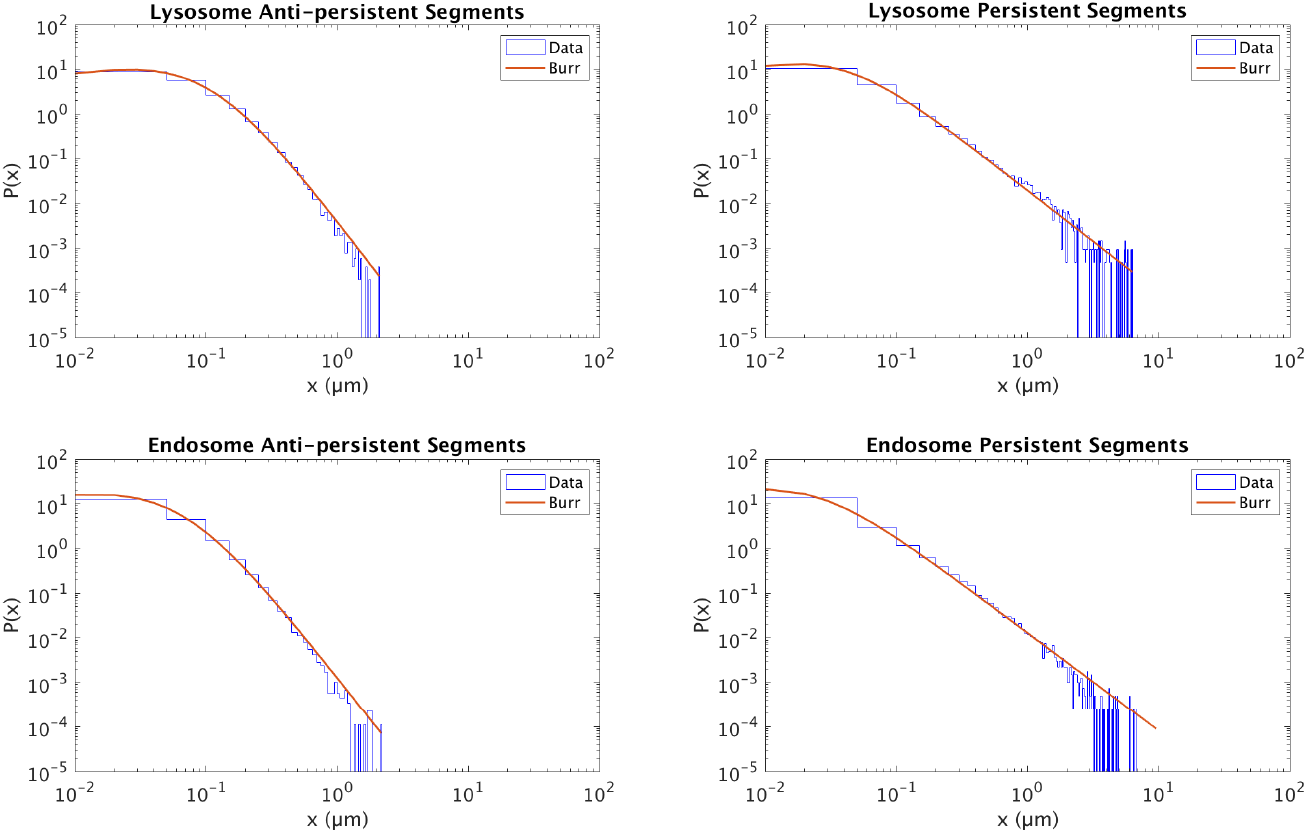
Normalized histograms (filled) and corresponding maximum likelihood estimation for Burr distributions (line) of segment displacements from lysosome and endosome experimental trajectories segmented using DLFNN. Parameter estimates are shown in Table 2.

